# Motor learning under mental fatigue: the compensatory role of rest periods

**DOI:** 10.64898/2026.03.21.713370

**Authors:** Célia Ruffino, Thomas Jacquet, Romuald Lepers, Charalambos Papaxanthis, Charlène Truong

## Abstract

Mental fatigue is known to impair cognitive and motor performance, but its impact on motor learning remains unclear. This study examined how mental fatigue affects skill acquisition in a sequential finger-tapping task. Twenty-eight participants were assigned to either a mental fatigue group, which completed a thirty-minute Stroop task, or a control group, which watched a documentary of equivalent duration. Both groups then trained on the finger-tapping task across multiple practice blocks with brief rest periods. Overall motor skill improved similarly in both groups. However, mental fatigue altered the pattern of acquisition: participants in the fatigue group showed decreased performance during practice blocks, which was compensated by larger gains during inter-block rest periods. A strong negative correlation was observed between online decrements and offline improvements, indicating that greater declines during practice were associated with larger gains during rest. This study highlights the critical role of rest periods in maintaining learning under cognitively demanding conditions and provides insight into how internal states, such as mental fatigue, can selectively influence the expression of performance without compromising overall learning.

## Introduction

Motor learning, defined as the process of acquiring, refining, and stabilizing new motor skills, is fundamental to human behavior and performance (Krakauer et al., 2019). It underpins performance in sport, the recovery of motor function during rehabilitation, and success in everyday activities ranging from simple tasks, such as walking or grasping, to complex actions, such as driving or playing a musical instrument. Given its essential role across these domains, understanding and optimizing motor learning is a major objective in both applied and fundamental neuroscience research.

During the acquisition phase of a motor skill, performance typically improves rapidly before reaching a plateau (Dayan & Cohen, 2011). Recent studies have shown that motor acquisition does not occur exclusively through practice, a process often referred to as online learning. Indeed, it also occurs during short rest periods interspersed within practice blocks. These so-called micro-offline processes support neural replay of trained movement sequences and contribute to performance gains during skill acquisition (Bönstrup et al., 2019, 2020). Additionally, these rest periods may help reduce reactive inhibition that accumulates during continuous practice and can otherwise lead to performance decrements (Rickard et al., 2008).

Motor learning does not solely result from practice structure. Learners’ internal states, such as rest or fatigue, may shape the balance between performance expression and learning-related processes. While early studies predominantly examined the effects of physical fatigue on motor learning (Alderman, 1965; Carron, 1969; Cotten et al., 1972; Goushe et al., 2025), much less is known about how mental fatigue, arising from prolonged cognitive engagement, influences motor learning processes. Mental fatigue, defined as a psychobiological state resulting from prolonged or intense cognitive activity, is increasingly common in modern life (Boksem & Tops, 2008). It can result from a long day of computer work, extended classes, prolonged driving, or smartphone use. It has been shown to impair sustained attention, decision-making, motor control and reaction time (Boksem et al., 2005; Guo et al., 2018; Jacquet et al., 2020, 2023; Lorist et al., 2005) and to reduce physical performance (Marcora et al., 2009; Pageaux & Lepers, 2018; Van Cutsem et al., 2017).

Neurophysiological evidence suggests that these effects are partly mediated by alterations in activity within frontal brain regions, particularly the anterior cingulate cortex (ACC), a region critically involved in cognitive control, error monitoring, and performance regulation (Boksem et al., 2005; Lorist et al., 2005). Notably, the ACC also plays a key role in motor learning. Functional neuroimaging studies have shown ACC activation during the acquisition of complex motor sequences, where error monitoring and action adjustment are vital (Aizenstein et al., 2004; Jueptner et al., 1997). This functional overlap suggests that mental fatigue, by altering ACC-dependent control processes, may disrupt the dynamics of motor skill acquisition. To date, empirical evidence addressing this issue remains scarce (Goushe et al., 2025). One study has reported a facilitating effect of mental fatigue during skill acquisition (Borragán et al., 2016). More recently, Godoi Filho et al. (2025) this relationship has been further nuanced by demonstrating that mental fatigue reliably impairs motor performance during skill acquisition. Together, these findings highlight the complexity of the relationship between mental fatigue and motor learning, and underscore the need to clarify its role in skill acquisition.

Therefore, the aim of this study was to examine the effect of mental fatigue on motor skill acquisition in a sequential finger-tapping task (F.T.T) across two groups of participants who trained either after a mentally fatiguing task or after a cognitively non-fatiguing control task. Based on evidence linking mental fatigue to impaired cognitive control and motor performance (Habay et al., 2026; Van Cutsem et al., 2017), and on the role of the ACC in motor learning (Aizenstein et al., 2004; Jueptner et al., 1997), we hypothesized that mental fatigue would alter the dynamics of motor skill acquisition.

## Method

### Participants

The sample size was determined based on previous studies using a similar experimental design, task, and outcome measures, which typically included 12–14 participants per group and were sufficient to detect meaningful changes in motor skill acquisition (Ruffino et al., 2021; Truong et al., 2023). Given the absence of a well-established smallest effect size of interest for the effect of mental fatigue on motor skill acquisition, no formal a priori power analysis was conducted. Instead, we report the sensitivity of the present design, which was adequate to detect medium effect sizes in between-group comparisons (approximately Cohen’s d ≈ 0.55), consistent with effect sizes commonly reported in the motor learning literature. This approach allows an informed interpretation of both significant and non-significant results.

Thus, twenty-eight healthy adults, free from neurological or physical disorders, participated in the present study after providing informed consent. According to the Edinburgh Handedness Questionnaire (Oldfield, 1971), all participants were right-handed (mean = 0.89 ± 0.13). Due to the nature of our fine SFTT, musicians and professional typists were excluded. Participants were advised to sleep for at least 8 hours (reported mean sleep duration was 8 ± 1 hours in both groups), abstain from alcohol, and avoid vigorous physical activity on the day before the visit. They were also advised against consuming caffeine or nicotine before the experiment. Participants were randomly assigned to two groups: the Mental Fatigue group (n = 14, 5 females, mean age: 25 ± 7 years) and the Control group (n = 13, 8 females, mean age: 28 ± 4 years). The experiment was conducted in conformity with the latest version of the Declaration of Helsinki (1964) and approved by the local Ethics Committee of the Université Bourgogne Franche-Comté (CERUBFC-2023-02-01-005).

### Procedure

#### General procedure

Participants attended the laboratory for a single experimental session (Figure 1), which was consistently scheduled for 9 a.m. After a brief familiarization phase, with the cognitively demanding task (Stroop task) for the Mental Fatigue group, and with the finger-tapping task (F.T.T) for both groups, participants completed psychological assessments, including subjective measures of mental fatigue and sleepiness. Then, participants of the Mental Fatigue group performed a 30-minute Stroop task, while those in the Control group watched an emotionally neutral documentary. Afterwards, all participants repeated the same psychological assessments (mental fatigue and sleepiness) and additional subjective measures of workload (NASA-TLX) and boredom. Subsequently, participants completed 48 trials of F.T.T (12 blocks of 4 trials, Truong et al., 2023) with 30-second rest periods between blocks to avoid mental fatigue (Rozand et al., 2016). Skill improvement was assessed by comparing the first two trials (trials 1 and 2, pre-test) with the last two trials (trials 47 and 48, post-test).

**Figure 1.**
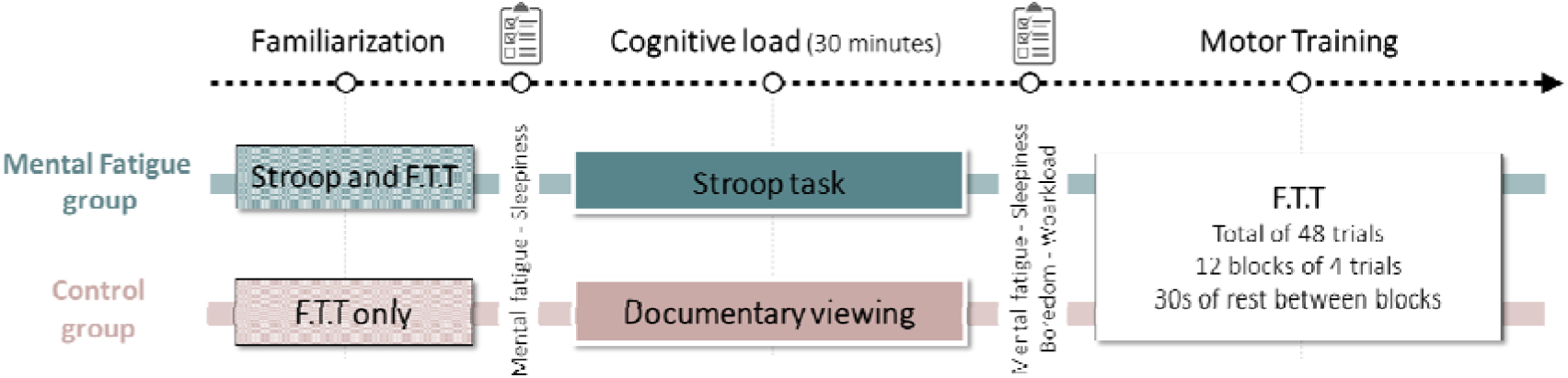
Experimental design and procedures. Participants in both the Mental Fatigue and Control groups began with a familiarization phase, followed by initial psychological assessments. Subsequently, participants were exposed to the cognitive load manipulation: a 30-minute Stroop task for the Mental Fatigue group or a neutral documentary viewing for the Control group. Second psychological assessments followed. Finally, participants completed motor training on the finger-tapping task (F.T.T), consisting of 12 blocks of 4 trials, with 30-second rest periods between blocks.

#### Cognitively demanding task

The cognitively demanding task was a modified Stroop task, previously used successfully to induce mental fatigue (Mangin et al., 2021). Each trial begins with the presentation of a fixation cross in the center of the screen, surrounded by either a square or a circle (50 ms). The fixation cross then remains displayed in the center of the screen for 400 ms. Next, a word appears at the center of the screen for 1250 ms, or until the participant responds. The words displayed (i.e., the response signals) are color names (red, blue, yellow, or green) written in ink of a different color (red, blue, yellow, or green). In this version of the Stroop task, the word is always incongruent with the ink color (e.g., the word “red” written in blue ink). When a square surround the fixation cross, participants are required to read the word, and when it is a circle, they must name the ink color. Participants have top of 1250 ms to respond, followed by a 300 ms fixation cross. Thus, each trial lasts 2 seconds. The task lasts 30 minutes and consists of 900 trials.

#### Documentary watching

The control task involves watching a documentary for 30 minutes, which matches the duration of the cognitively demanding task. Participants will choose between two documentaries: ‘Legacy’ by Y.A. Bertrand or ‘Bill Gates’ by S. Malterre.

#### Finger-tapping task (F.T.T)

We used a computerized version of the sequential finger-tapping task (Kami et al., 1995), which is commonly employed in laboratory experiments (Truong et al., 2023, 2024). Participants were comfortably seated on a chair in front of a keyboard. They were requested to tap, as accurately and as fast as possible, with their left hand (non-dominant hand) the following sequence: 3-1-4-2-3-0. Each key was assigned to a specific finger: 0 (thumb), 1 (index), 2 (middle), 3 (ring), and 4 (little). One trial consisted of six sequences. Specifically, at the beginning of each trial, participants pressed the key ‘0’ with their thumb to start the chronometer, and they accomplished the 6 sequences continuously. Pressing the key ‘0’ at the end of the 6th sequence stopped the chronometer and ended the trial. The view of the non-dominant hand was occluded by a box throughout the protocol. The sequence’s order was displayed on the box and visible to participants throughout the experiment.

#### Psychological assessments

##### Mental fatigue level

Subjective mental fatigue was assessed using a visual analogue scale (VAS) (Smith et al., 2019), administered before and after the cognitively demanding task, as well as prior to and following the documentary viewing. The VAS consisted of a 100-mm line with bipolar endpoints (0 mm = ‘Not tired at all’; 100 mm = ‘Extremely tired’). Participants were asked, ‘What is your current level of mental fatigue?’ and instructed to place a mark on the line to indicate their level of fatigue. The score was calculated by measuring the distance from the left anchor (0 mm: ‘Not tired at all’) to the participant’s chosen mark.

##### Sleepiness level

Another visual analogue scale (VAS) was used to assess feelings of sleepiness (Alqurashi et al., 2021) before and after the cognitively demanding task and the documentary viewing. This VAS consisted of a 100-mm line with bipolar end points (0 mm = ‘Not sleepy at all’; 100 mm = ‘Extremely sleepy’). To measure sleepiness, participants were asked ‘What is your current level of sleepiness?’ and instructed to place a mark along the line to indicate their feelings. The VAS score was calculated by measuring the distance from the left anchor (0 mm: ‘Not sleepy at all’) to the participant’s chosen mark.

##### Boredom level

Feelings of boredom following the cognitively demanding task and the documentary viewing were assessed using a Visual Analogue Scale (VAS) with endpoints ranging from 0 mm (“Not bored at all”) to 100 mm (“Extremely bored”). Participants were prompted with the question, “How bored do you feel right now?” and instructed to mark their current level of boredom on the scale. The score was calculated by measuring the distance from the starting point (0 mm) to the participant’s mark.

##### Subjective workload

The National Aeronautics and Space Administration Task Load Index (NASA-TLX) was used to assess subjective workload (Hart & Staveland, 1988). The NASA-TLX consists of six subscales: mental demand (how much mental and perceptual activity was required?), physical demand (how much physical activity was required?), temporal demand (how much time pressure did you feel due to the pace or rate at which the task occurred?), performance (how successful do you think you were in accomplishing the goals set by the experimenter?), effort (how hard did you have to work to achieve your level of performance?), and frustration (how irritating or annoying did you find the task?). Participants rated each item on a scale divided into 20 equal intervals, anchored by bipolar descriptors (e.g., high/low). The score for each subscale was then multiplied by 5, yielding a final score between 0 and 100 for each subscale.

### Data recording and analysis

For the F.T.T, movement accuracy and duration were calculated for each trial. Accuracy, referred to as the ‘Error rate,’ was defined as the number of incorrect sequences within a trial (0 = no errors; 6 = all sequences incorrect). A trial was considered incorrect if the participant made one or more mistakes in the sequence. The error rate was determined as the percentage of errors relative to the total number of possible errors within the trial:

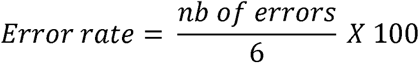

Movement duration was defined as the time elapsed (in seconds) from the start of the trial (when the participant pressed the ‘0’ key) to its end (when the participant pressed the ‘0’ key after completing the 6th sequence).

These two parameters (Movement duration and Error rate) are linked through the speed-accuracy tradeoff function (Shmuelof et al., 2012). Therefore, motor skill improvement (i.e., the training-induced change in the speed-accuracy tradeoff) cannot occur when duration and accuracy change by the same magnitude in opposite directions. To address this, we calculate a composite ratio (in arbitrary units, a.u.) to characterize motor skill as follows (Ruffino et al., 2021; Truong et al., 2023, 2024) :

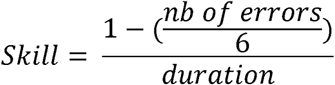

In that formula, skill increases if the duration decreases and/or if the number of errors decreases.

Gains between pre- and post-test were calculated following a simple proportional formula that indicates the amount of skill acquisition:

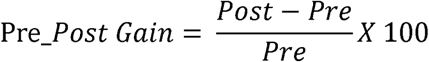

Online changes in motor acquisition were determined by subtracting the first trial from the last trial within each block and then summing these values across all blocks. Similarly, offline changes in motor acquisition were computed by subtracting the last trial of the previous block from the first trial of the current block, and summing the results. Finally, the net change was obtained by adding the online and offline changes (see Figure 3C).

### Statistical analysis

One participant was excluded from the statistical analyses because of a negative acquisition gain, indicating the absence of learning. Consequently, the final sample comprised 27 participants (Mental Fatigue group: n = 14; Control group: n = 13). Data normality was evaluated with the Shapiro-Wilk test, while sphericity was examined using Mauchly’s test. Suitable statistical tests were chosen based on these assumptions. For all analyses, the significance level was set at 0.05, and the observed power exceeded 0.8.

Subjective measures of mental fatigue and sleepiness were analyzed using a repeated-measures (rm) ANOVA, with *Group* (Mental Fatigue and Control group) as a between-subjects factor and *Time* (before and after the cognitively demanding task) as a within-subjects factor. Bonferroni post hoc tests were applied when necessary. Between-group comparisons of subjective boredom and workload (recorded after the cognitively demanding task) were analyzed using Mann–Whitney *U* tests due to non-normality.

Motor skill acquisition in F.T.T was assessed using rm ANOVA with *Group* (Mental Fatigue group and Control group) as between-subjects factor and *Test* (Pre-and Post-Test) as within-subjects factor. Bonferroni post hoc tests were applied. Additionally, the gain in motor skill (Pre_Post gain) was compared between groups using an independent t-test, and each group’s gain was compared with the reference value of zero (*t-test*).

Identical analyses were conducted for the movement duration (see Supplementary Result 1). Non-normal error rate distributions (Shapiro-Wilk test, *p* < 0.05) were analyzed using two-tailed permutation tests (5000 permutations). Multiple comparisons were corrected using the Benjamini-Hochberg False Discovery Rate (see Supplementary Result 1).

The improvement of individual skill performance across the 48 training trials was modeled using a power function:

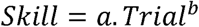

where a represents the initial amplitude of the learning process, and b denotes the learning rate. Between-group differences in learning rate (b) were analyzed using Mann–Whitney *U* tests.

To further characterize the shape of each participant’s learning curve, we identified the first trial of the performance plateau. Precisely, this was defined as the trial at which the derivative of the fitted power function (i.e., the rate of change in skill between consecutive trials) dropped to within 0.0001 above its minimum. This time point was compared between groups using a Mann–Whitney *U* test. To assess the robustness of this criterion, the same analysis was repeated using alternative thresholds (0.0002, 0.0003, 0.0004, and 0.0005; see Supplementary Result 2).

Depending on the normality assumption, differences between groups in online, offline, and net changes were assessed using independent t-tests or Mann–Whitney *U* tests. Each change score was also compared to the reference value of zero (*t-test*). Additionally, Pearson correlation coefficients were calculated to examine the relationship between online and offline changes.

To strengthen conclusions in cases of non-significant results, Bayesian equivalence analysis was conducted with a region of practical equivalence (ROPE) = [−0.1,0.1] and a prior Cauchy scale of 0.707 (Morey & Rouder, 2011). This analysis quantifies the evidence in favor of the null hypothesis and discriminates the ‘absence of evidence’ and the ‘evidence of absence’, leading to stronger and more reliable conclusions.

Effect sizes were reported as partial eta squared (η*^2^_p_*) for ANOVA, categorized as small (≥ 0.01), moderate (≥ 0.07), or large (≥ 0.14). For t-tests, Cohen’s d was reported with small (≥ 0.20), moderate (≥ 0.50), and large (≥ 0.80) effects. Rank-biserial correlations (*rb*) were reported for non-parametric (Mann–Whitney *U*) comparisons with small (≥ 0.10), moderate (≥ 0.30), and large (≥ 0.50) effects.

## Results

### Cognitive demanding task: psychological assessment

Figure 2A illustrates the average subjective feeling of mental fatigue. A repeated measures ANOVA revealed a significant main effect of *time* (*F*_1, 25_ = 56.63, *p* < .001, η*^2^_p_* = 0.69) and *group* (*F*_1, 25_ = 4.33, *p* < 0.048, η*^2^_p_* = 0.15), as well as a significant *time x group* interaction (*F*_1, 25_ = 37.49, *p* < .001, η*^2^_p_* = 0.60). Post hoc analyses showed no difference in subjective mental fatigue between groups before the cognitively demanding task or the documentary viewing (*p* = 1.00). However, a significant difference emerged in the Mental Fatigue Group after the tasks (*p* < 0.001). Moreover, the Mental Fatigue group exhibited a significant increase in fatigue from pre- to post-test (*p* < 0.001), whereas no change was observed in the Control group (*p* = 1.00).

**Figure 2.**
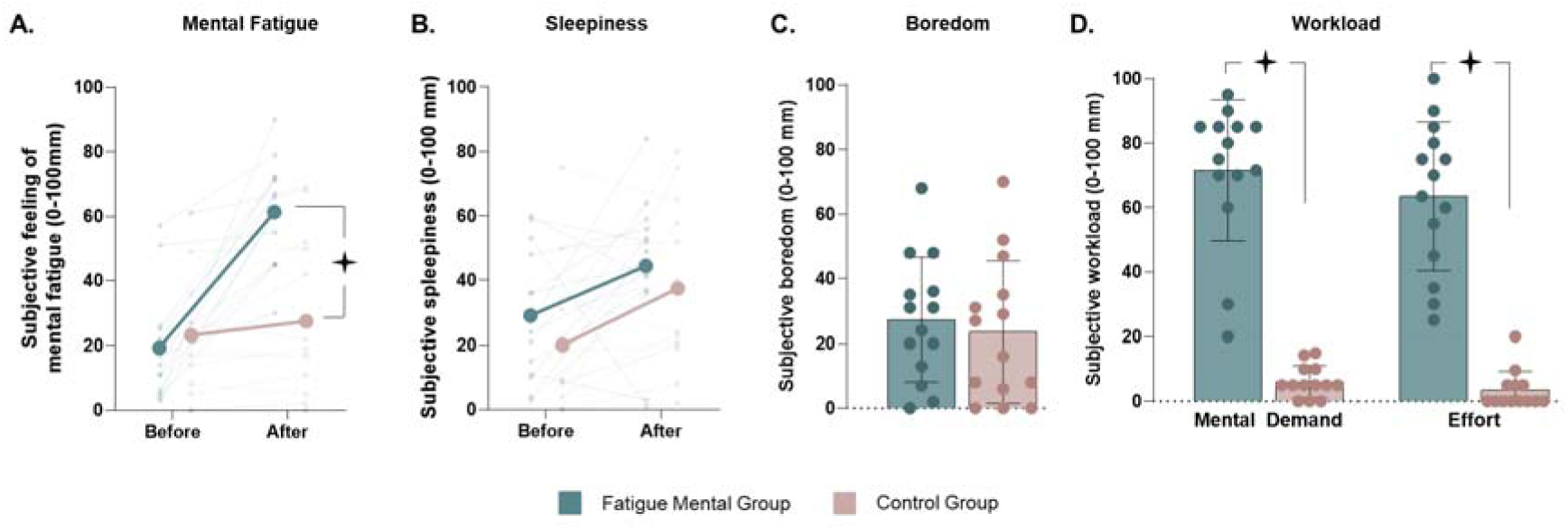
Psychological assessment of Mental Fatigue Group and Control Group. **A** - **B.** Average values of subjective feeling of fatigue and sleepiness respectively before and after Stroop task (Mental Fatigue Group) or documentary watching (Control Group). **C** - **D.** Average values (+SD) of subjective boredom and subjective workload (NASA-TLX), respectively, after Stroop task (Mental Fatigue group) or documentary watching (Control Group). Dots represent individual data. Black stars indicate a difference between groups.

Regarding subjective sleepiness (Figure 2B), we found a significant main effect of *time* (*F*_1, 25_ = 9.16, *p* = 0.006, η*^2^_p_* = 0.27), indicating an overall increase in sleepiness regardless of the task but no significant group *effect* (*F*_1, 25_ = 1.39, *p* = 0.25, η*^2^_p_* = 0.05) or interaction (*F*_1, 25_ = .04, *p* = 0.85, η*^2^_p_* = 0.002). Similarly, subjective boredom (Figure 2C) did not differ significantly between the cognitively demanding task and the documentary (*U* = 74.50, *p* = 0.87, *rb* = −0.05).

Concerning subjective workload, analyses of the NASA-TLX (Figure 2D) showed that the cognitively demanding task was perceived as significantly more mentally (*U* = 0.00, *p* < 0.001, *rb* = −1.00), temporally (*U* = 2.00, *p* < 0.001, *rb* = −0.98) demanding, as well as requiring more effort (*U* = 0.00, *p* < 0.001, *rb* = −1.00) and causing greater frustration (*U* = 2.50, *p* < 0.001, *rb* = −0.97) than watching the documentary. Only physical demanding did not differ significantly between the two groups (*U* = 60.00, *p* = 0.08, *rb* = −0.34).

Taken together, these results confirm that the cognitively demanding task induces significant mental fatigue without affecting sleepiness or boredom.

### Motor skill acquisition

Figure 3A shows the motor skill at Pre and Post for the Control and Mental Fatigue Group. A repeated-measures ANOVA revealed a significant main effect of *Test* (*F*_1, 25_ = 149.24, *p <* .001, η*^2^_p_* = 0.86), indicating a strong improvement in motor skill between pre- and post-test. However, no significant interaction was found (*F*_1, 25_ = 0.37, *p =* 0.55, η*^2^_p_* = 0.02), nor a main effect of *group* (*F*_1, 26_ = 0.84, *p =* 0.37, η*^2^_p_* = 0.03), suggesting that the observed improvement occurred regardless of the group.

**Figure 3.**
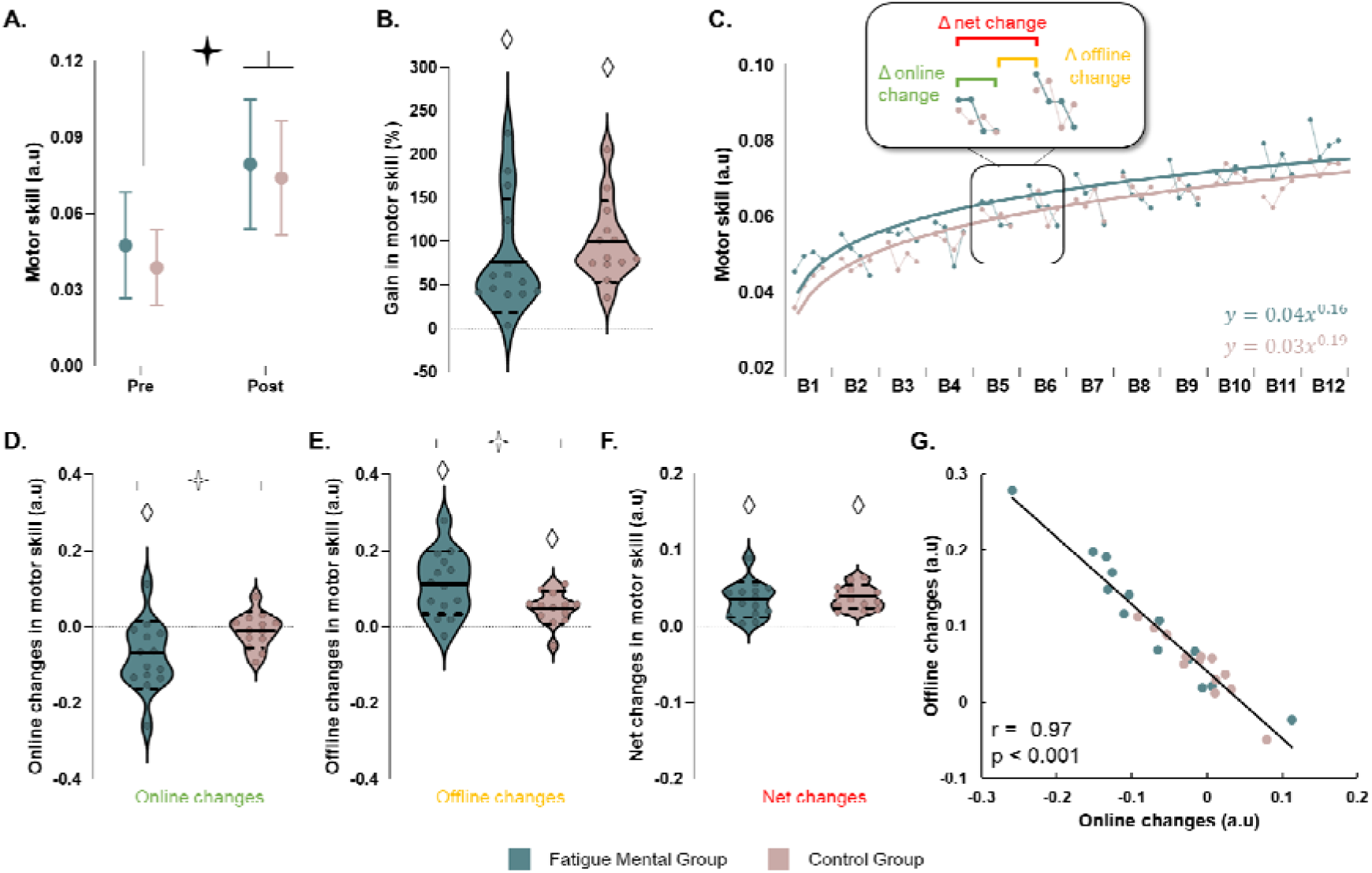
Acquisition analysis. **A.** Average values (+SD) of motor skill in Pre- and Post-Test for each group.. **B.** Violin plots for the percentage of Pre_Post gain in motor skill for each group. Dots represent individual data. Thick and thin horizontal lines mark mean and SD, respectively. **C.** Trial-by-trial plotting of motor skill during each block (B1-B12) of the training. Dots correspond to the group means for each trial and curves correspond to the power law functions using the mean point for each trial. Box represents a zoom on B5 and B6 and shows the online, offline and net change between their both blocks. **D-E-F.** Violin plots for the online, offline and net changes, respectively, for each group. Dots represent individual data. Thick and thin horizontal lines mark mean and SD, respectively. **G.** Correlation between offline changes and online changes. Black star indicates a significant difference between Pre- and Post-test. White stars indicate a significant different between groups. White diamonds indicate significant differences from the value zero.

Similarly, the comparison of the Pre_Post gains with the reference value *zero (*Figure 3B) confirmed significant improvement for the Mental Fatigue group (*t =* 4.78, *p <* 0.001, *d =* 1.29) and for the Control group (*t =* 7.89, *p <* 0.001, *d =* 2.19) without difference between both groups (*t =* 0.79, *p =* 0.44, *d =* 0.30. Complementary, Bayesian equivalence testing on the Pre_Post gains between groups showed that the data were 2.2 times more likely to validate the null hypothesis than the alternative one (BF_OH_^01^) and to lie in the equivalence than in the non-equivalence region (BF_NOH_^01^).

To further explore acquisition, we examined the skill learning curves adjusted using a power function (Figure 3C). The analysis did not reveal a significant difference between groups (factor b: *U* < 120, *p =* 0.17, *rb* = 0.32; Bayesian equivalence test: BF_OH_^01^ = 2.19; BF_NOH_^01^ = 2.21), indicating that the learning rate did not differ according to the task performed before F.T.T. Furthermore, we found no variation in the point at which the skill improvement plateaued (Mental Fatigue group: 27□±□6 trials; Control group: 30□±□4 trials; *U* = 117.50, *p* = 0.21, *rb* = 0.29; Bayesian equivalence tests: BF_OH_^01^ = 1.51; BF_NOH_^01^ = 1.52), regardless of the threshold selected (see Supplementary Result 2).

Interestingly, the Mental Fatigue group exhibited a greater decline in online changes of motor skill through block trials (i.e., the sum for each block of the change between the first and the last trial of the block), relative to the Control group (*t* = 2.41, *p* = 0.02, *d* = 0.93, Figure 3D). Comparison with the reference value *zero* indicated significant negative changes in motor skill for the Mental Fatigue Group (*t* = −3.23, *p* < 0.001, *d* = −0.86) but not for the Control Group (*t* = −0.83, *p* = 0.42, *d* = −0.23; Bayesian equivalence tests: BF_OH_^01^ = 2.67; BF_NOH_^01^ = 2.71).

The opposite pattern was observed for the offline changes (Figure 3E, that is, the total of the change between the last trial and the first trial of the next block during each inter-block rest period), with a greater increase in motor skill for the Mental Fatigue group than the Control group (*U* = 45, *p* = 0.03, *rb* = 0.22). Comparison with the reference value *zero* indicated a significant positive change in motor skill for both groups (*t* > 4.18, *p* < 0.001, *d* > 1.16 for all).

Analysis of the net changes (i.e., the sum of the online and offline changes) revealed that it was significantly positive in both groups (comparison with zero: *t > 5.57*, *p <* 0.001, *d >* 1.49 for all, Figure 3F), indicating that the performance decline during practice was fully recovered during the inter-block rest periods, with additional improvement occurring. These net changes did not differ significantly between groups (*t* = 0.46, *p* = 0.65, *d* = 0.18; Bayesian equivalence tests: BF_OH_^01^ = 2.58; BF_NOH_^01^ = 2.62), supporting comparable overall learning.

Moreover, a strong negative correlation (*r* = –0.97, *p* < 0.001; Figure 3G) indicated that greater online declines were associated with greater offline gains, supporting the notion that declines were compensated during the inter-block rest periods.

Overall, the results indicate that although the mental fatigue group experienced a greater decrease during the online practice block, this decrease is compensated during inter-block rest periods, and finally, the skill improvement during the training sessions does not differ between groups, suggesting that the acquisition process is not impacted by the presence of mental fatigue, but is maintained by employing a compensatory strategy.

## Discussion

The present study aimed to investigate the impact of experimentally induced mental fatigue on motor skill acquisition using a sequential finger tapping task. Our results indicate that mental fatigue did not affect the overall amount of skill acquired by the end of practice, yet it substantially altered the temporal dynamics of its acquisition. Specifically, participants in the mentally fatigued condition exhibited reduced online performance gains within practice blocks, which were compensated for by offline improvements during rest periods. As a result, final performance levels were comparable between groups, despite markedly different acquisition profiles.

### Online performance decline under mental fatigue

Consistent with previous studies, repetition increases motor skill until a performance plateau is reached (Truong et al., 2023, 2024), regardless of cognitive fatigue state (Borragán et al., 2016; Godoi Filho et al., 2025). Mental fatigue, however, altered temporal dynamics: participants in the mentally fatigued group showed a significant decline in performance within practice blocks (online gains), while the control group maintained stable performance. Previous research on motor sequence learning suggests that such online performance decrements can arise from supraspinal mechanisms rather than peripheral fatigue, as markers of muscular fatigue often remain unchanged despite performance decline (Arias et al., 2015; Madrid et al., 2016; Rodrigues et al., 2009). For instance, breakdown of surround inhibition in primary motor cortex and increased coactivation of antagonistic muscles have been linked to transient performance drops followed by recovery during rest (Bächinger et al., 2019). A plausible hypothesis is that mental fatigue may exacerbate these mechanisms, leading to greater loss of online performance. Another possibility is that mental fatigue negatively affects performance expression, but this effect may be partially compensated at the beginning of each practice block, i.e., after a short rest. Indeed, it is well established that mental fatigue negatively affects motor performance, such as the speed-accuracy trade-off (Jacquet et al., 2020; Pageaux & Lepers, 2018; Rozand et al., 2016). In addition, recent work indicates that individuals can counteract mental fatigue, such as by increasing the effort invested in the task (Goushe et al., 2025; Wright & Mlynski, 2019), suggesting that performance can be temporarily maintained through enhanced cognitive resources. In this context, it is plausible that participants initially upregulate effort to sustain performance, but that this compensatory effort is limited and cannot be maintained over time.

#### Compensation via rest and explicit strategies

The observed decline in online performance within practice blocks was counteracted during inter-block rest periods, yielding micro-offline gains. The larger micro-offline gains in the mental fatigue group compared to control seem to be mainly attributable to reduced reactive inhibition that accumulates during continuous practice during rest periods (Rickard et al., 2008), as a strong negative correlation was observed between online decrements and offline improvements. Furthermore, in our study, participants could partially compensate for fatigue by using explicit strategies, thanks to the repetitive nature of the finger-tapping sequence. This was not possible in previous work by Godoi Filho et al. (2025), where the visuomotor tracking task used lacked repeating speed or amplitude patterns, preventing reliance on sequence-based strategies. Micro-offline gains may also include contributions from rapid consolidation processes, previously reported in motor sequence learning paradigms (Bönstrup et al., 2019; King et al., 2017). In our task, these contributions could not be precisely quantified due to early within-block performance drops. Trial duration differences (mean Pre-test duration: 23□±□8□s) may have further accelerated the observed performance decline compared to shorter trials used in other studies (Rickard et al., 2008; Robertson et al., 2004).

#### Final motor skill acquisition level is unaffected under mental fatigue

Despite transient online decrements, both groups reached comparable overall gains by the end of the training session. This suggests that mental fatigue primarily modulates performance expression rather than the underlying motor skill acquisition. This result diverges from Borragán et al. (2016), who reported a facilitative effect of mental fatigue on implicit motor learning. In that study, mental fatigue was proposed to reduce the availability of attentional and executive resources, thereby limiting engagement of the explicit learning system and promoting relative reliance on implicit learning mechanisms, consistent with previous theoretical accounts (Brown & Robertson, 2007; Robertson et al., 2004).

This interpretation is further supported by evidence that mental fatigue disproportionately affects prefrontal–cingulate networks, particularly the dorsolateral prefrontal cortex (DLPFC) and ACC (Wylie et al., 2020; Yan et al., 2025). These regions are critically involved in attentional control, working memory, performance monitoring, and the implementation of explicit strategies during motor learning (Carter et al., 1998; Lin et al., 2022; Nguyen et al., 2025). In the present study, participants were required to memorize and reproduce a predefined numerical sequence, a process strongly dependent on these explicit mechanisms. Because explicit processes cannot be bypassed in this task, unlike in implicit tasks, which can often be performed automatically (Borragán et al., 2016), mental fatigue did not facilitate acquisition, which may explain the discrepancy with previous findings.

#### Conclusion and future directions

The present study demonstrates that mental fatigue reshapes the temporal dynamics of motor performance during practice, leading to transient decrements without affecting overall skill acquisition. Our results highlight the critical role of rest periods in restoring performance under mental fatigue, suggesting that short-term recovery mechanisms help maintain overall learning despite temporary performance drops. Importantly, the current study primarily investigated fast processes of motor learning associated with skill acquisition, but future work should also examine slower consolidation processes, which are crucial for long-term retention. If mental fatigue reduces engagement of explicit processes, this reduction could potentially facilitate consolidation. Indeed, competition between motor and declarative memory systems during learning can limit consolidation, and reducing this competition, for example, by perturbing declarative processes immediately after acquisition, has been shown to stabilize motor memory (Breton & Robertson, 2014; Truong et al., 2024; Tunovic et al., 2014). Future studies combining behavioral and neurophysiological measures would help clarify the neural mechanisms through which mental fatigue influences these performance dynamics, particularly through interactions between executive control structures (e.g., DLPFC and ACC) and the mechanisms underlying performance decrement and recovery. A better understanding of these interactions could inform training and rehabilitation protocols by helping optimize skill acquisition under conditions of mental fatigue.

## Supporting information

Supplementary Result

## Data availability

All data from this study are available at: https://osf.io/vh3c7/overview?view_only=18c8afb9d6504a8f8bfbc6fc4c5ecb74

## Author contributions

CR and TJ designed the experiment and recorded the data; CT analyzed the data and developed figures; CR and CT wrote the manuscript; TJ, RL, and CP provided feedback on the manuscript; all co-authors read and approved the submitted version.

