## Supplementary Result for "Motor learning under mental fatigue: the compensatory role of rest periods"

### Supplementary Information

**Supplementary Result 1:** Movement duration and Error rates

We conducted a repeated-measures (rm) ANOVA on movement duration, with group (as the between-subjects factor and session (Pre, Post) as the within-subjects factor. Post hoc comparisons were carried out using Bonferroni tests. In addition, because the distribution of error rates deviated from normality, we performed non-parametric permutation tests with Benjamini–Hochberg correction. Supplementary Figure S1 illustrates the mean values (+SD) of movement duration and error rates for Mental Fatigue and Control group. Skill improvement from Pre to Post was mainly explained by faster movements (only session effect was found: *F*_1,25_ = 107.38, *p* < 0.001, *ηp^2^* = 0.81) while no significant effect was found for accuracy (-0.38 < *T* < 1.66, p > 0.43 in all cases).

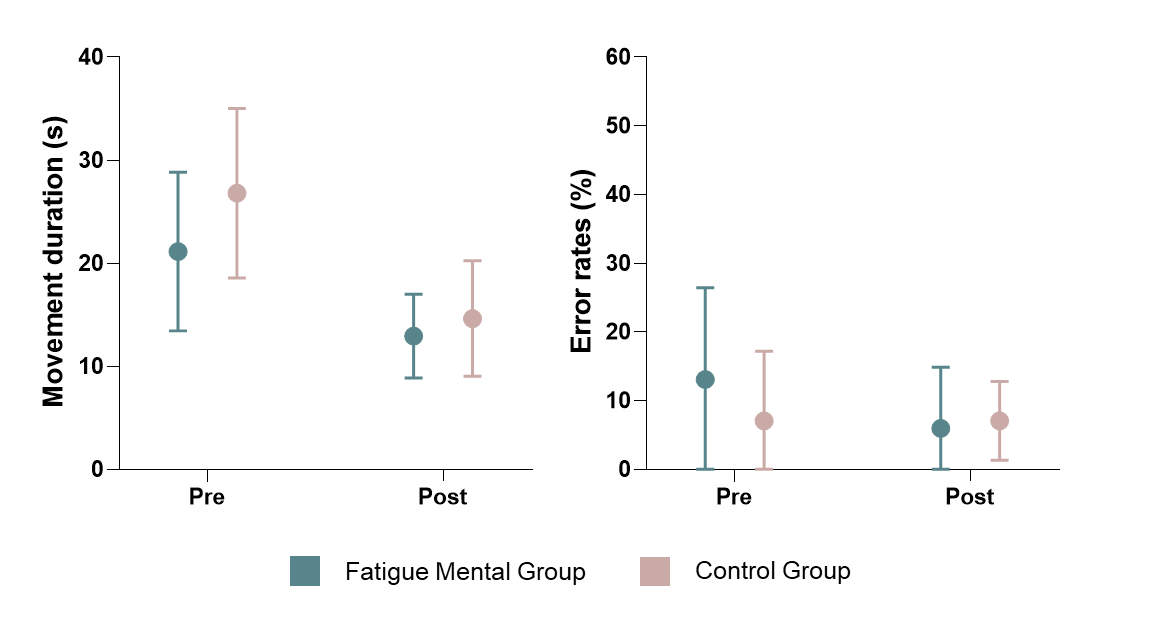

**Supplementary Figure S1.** Mean (+SD) of movement duration (left) and error rates (right) in Pre- and Post- tests for (a) Fatigue Mental and Control groups. The star (⯎) indicates significant differences between sessions.

**Supplementary Result 2:** Threshold values for the plateau performance

Since our threshold value of 0.001 was arbitrarily determined, we performed, as a control, the same analysis with a threshold value of 0.0002, 0.0003, 0.0004, and 0.0005. Supplemental Table S1 indicates the mean (+SD) of the trial when groups reached the performance plateau according to the five different threshold values. Independent t-tests were performed to compare group plateaus at each threshold. Results indicated no significant differences between groups regardless of the threshold applied.

**Tableau S1.** Supplementary Results. The mean (+SD) of the first trial of the performance plateau according to the different threshold values for each group, including *t* and *p* values for between-group comparisons.

| **Experiment 1** | | | | |  |  |
| --- | --- | --- | --- | --- | --- | --- |
|  |  | **0.0001** | **0.0002** | **0.0003** | **0.0004** | **0.0005** |
| **Fatigue Mental** | *Mean* | 27.43 | 19.64 | 15 | 12.21 | 10.14 |
|  | *SD* | *6.49* | *6.54* | *6.08* | *5.25* | *4.62* |
| **Control** | *Mean* | 30.23 | 22.23 | 17.46 | 13.92 | 11.77 |
|  | *SD* | *4.49* | *4.80* | *4.31* | *3.73* | *3.47* |
|  | *Stat (t)* | 1.29 | 1.17 | 1.21 | 0.97 | 1.03 |
|  | ***p-value*** | **0.21** | **0.26** | **0.24** | **0.34** | **0.31** |
